# Effect of psilocybin on marble-burying in ICR mice: Role of 5-HT1A receptors and implications for the treatment of obsessive-compulsive disorder

**DOI:** 10.1101/2022.07.13.498401

**Authors:** Sandeep Singh, Alexander Botvinnik, Orr Shahar, Gilly Wolf, Corel Yakobi, Michal Saban, Adham Salama, Amit Lotan, Bernard Lerer, Tzuri Lifschytz

## Abstract

**Background:** Preliminary clinical findings, supported by preclinical studies employing behavioral paradigms such as marble-burying, suggest that psilocybin may be effective in treating obsessive-compulsive disorder.

**Aims:** To explore 1) the role of 5-HT2A and 5-HT1A receptors in the effect of psilocybin on marble-burying; 2) the effect of staggered versus bolus psilocybin administration and persistence of the effect; 3) the effect of the 5-HT1A partial agonist, buspirone, on marble-burying and the head-twitch response (HTR) induced by psilocybin, a rodent correlate of psychedelic effects.

**Methods:** Male ICR mice were administered psilocybin 4.4 mg/kg, escitalopram 5 mg/kg, 8-OH-DPAT 2 mg/kg, M100907 2 mg/kg, buspirone 5 mg/kg, WAY100635 2 mg/kg or combinations, intraperitoneally, and were tested on the MBT. HTR was examined in a magnetometer-based assay.

**Results:** 1) Psilocybin and escitalopram significantly reduced marble-burying. The effect of psilocybin was not attenuated by the 5-HT2A antagonist, M100907. The 5-HT1A agonist, 8-OH-DPAT reduced marble-burying as did the 5-HT1A partial agonist, buspirone. The effect of 8-OH-DPAT was additive to that of psilocybin but that of buspirone was not. The 5-HT1A antagonist, WAY100635, attenuated the effect of 8-OH-DPAT and buspirone but not the effect of psilocybin. 2) Psilocybin injections over 3.5 hours had no effect on marble-burying and the effect of bolus injection was not persistent. 3) Co-administration of buspirone with psilocybin blocked its effect on HTR

**Conclusions:** Neither 5-HT2A nor 5-HT1A receptors are pivotally implicated in the effect of psilocybin on marble-burying. Co-administration with buspirone may block the psychedelic effects of psilocybin without impeding its anti-obsessional effects.

## Introduction

Obsessive compulsive disorder (OCD) is a serious, often disabling disorder that affects 100-150 million people with an estimated worldwide prevalence of up to 2% (Sasson et al, 1997). It is characterized by uncontrollable, recurring thoughts (obsessions) that cause significant distress and by behaviors and actions that the sufferer has the urge to repeat over and over. Available treatments include tricyclic antidepressants such as clomipramine that block serotonin reuptake, specific serotonin reuptake inhibitors (SSRIs), augmentation with second generation antipsychotic drugs that are 5-HT2A receptor blockers and exposure and response prevention, a form of cognitive behavioral therapy (Marek et al., 2005). At least a third of patients with OCD do not respond to first-line treatments (Bloch et al., 2013). There is a clear need for therapeutic alternatives.

Research on the clinical application of psychedelics in psychiatry is growing apace after several decades in which their use was legally proscribed (Nutt, 2019; Rucker and Young, 2021). These drugs, particularly the classical tryptaminergic psychedelics, act principally but not only via the 5-HT2A receptor to induce profound changes in perception, cognition, and mood (Nichols, 2016; Rucker et al., 2016). There is intriguing but limited preliminary evidence that psychedelics, specifically psilocybin, may have a role in the treatment of OCD. This includes a case report on the effectiveness of self-medication with psilocybin-containing magic mushrooms in a patient with long-standing OCD (Lugo-Radillo and Cortes-Lopez, 2021). A preliminary clinical trial of a range of psilocybin doses in 9 patients yielded positive results which were not necessarily related to the induction of psychedelic effects (Moreno et al, 2006). There are no published reports since the Moreno et al (2006) study but several clinical trials of psilocybin in OCD and related disorders are currently underway (https://www.clinicaltrials.gov/ct2/results?cond=psilocybin&term=ocd&cntry=&state=&city=&dist=, Accessed June 27, 2022)..

In the preclinical context, the marble-burying test (MBT) uses the propensity of rodents to dig and cover non-threatening objects placed in their cages as a predictor of anti-obsessional effects (Thomas et al., 2009;). Although criticized as a model of OCD (de Brouwer et al., 2019), MBT has reasonable construct validity in terms of identifying drugs that are effective in OCD and potential treatments for the disorder (Korff and Harvey, 2006). Studies with MBT have yielded results supportive of a potential therapeutic effect of psychedelics in OCD. Two studies have shown an effect of psilocybin to significantly reduce marble-burying in mice (Matsushima et al., 2009; Odland et al., 2021a). Similar results have been reported for 2,5-dimethoxy-4-iodoamphetamine (DOI) (Njung’e and Handley, 1991; Egashira et al., 2012; Odland et al., 2021a, 2021b) and for the highly selective, brain-penetrant 5-HT2A receptor agonist N-(2-hydroxybenzyl)-2, 5-dimethoxy-4-cyanophenylethylamine (25CN-NBOH) (Odland et al., 2021b).

Psilocin, the active metabolite of psilocybin, binds to 5-HT2A receptors with high affinity (Madsen et al., 2019). Psychedelic effects of psilocybin in humans are blocked by the 5-HT2A receptor antagonist, ketanserin (Quednow et al., 2012), as is the effect of psilocybin to induce the head twitch response (HTR) in mice (Halberstadt and Geyer, 2013; Klein et al., 2021), a characteristic rodent behavior that is correlated with psychedelic effects in humans (Halberstadt et al., 2020). The anti-marble-burying effect of psilocybin does not appear to be mediated by 5-HT2A receptors since it is not blocked by prior administration of the highly selective 5-HT2A antagonist, M100907 (volinanserin) (Odland et al, 2021a). It is noteworthy, however, that M100907 does block the effect of DOI to reduce marble-burying (Odland, et al., 2021a). Odland et al (2021a) also showed that the effect of psilocybin on marble-burying was not blocked by the 5-HT2C receptor antagonist, SB242084, suggesting that this receptor is also not implicated in the anti-marble-burying effect of psilocybin.

In the current study, we sought to sought to re-examine the role of 5-HT2A receptors in the effect of psilocybin on marble-burying and to explore the role of 5-HT1A receptors by studying 5-HT1A agonist, partial agonist and antagonist effects in the MBT in conjunction with psilocybin. We also examined the effect of staggered psilocybin administration as compared to bolus injection and persistence of the anti-marble-burying effect after 7 days.

## Materials and Methods

### Ethical Statement

Experiments were conducted in accordance with AAALAC guidelines and were approved by the Authority for Biological and Biomedical Models Hebrew University of Jerusalem, Israel, Animal Care and Use Committee (Project No. MD-21-16663-4 approved on 22/8/2021 and Project No. MD-21-16754-4 approved on 12/10/2021). All efforts were made to minimize animal suffering and the number of animals used.

### Reagents and chemicals

Chemically synthesized psilocybin (PSIL) was kindly supplied by USONA Institute, Madison, Wisconsin, USA and was determined by Ultra Performance Liquid Chromatography (UPLC) to contain 98.75% psilocybin. Escitalopram (ESC), 8-hydroxy-dipropylamino-tetralin hydrobromide (8-OH-DPAT) (DPAT), volinanserin (M100907) and buspirone (BUSP) were of highest purity grade and were purchased from Sigma Aldrich Israel Ltd, Israel. WAY100635 (WAY) was obtained from Biotest (Tocris Bioscience), Israel. 8-OH-DPAT and M100907 were dissolved in vehicle (0.9 % saline) (VEH) containing 5% dimethyl sulfoxide (DMSO). Psilocybin, escitalopram, WAY100635 and buspirone were dissolved in 0.9% saline vehicle.

### Animals and Experimental design

The study was carried out on male ICR mice (30.0 ± 2.0 gm). The animals were housed in a controlled environment (25 ± 2 °C, relative humidity 55 ± 15 %) with a 12-h light/dark cycle. All mice were fed with a normal laboratory diet of nutrient-rich pellets ad libitum. The experimental mice were allowed to acclimatize prior to starting the experiment.

Drugs were administered by intraperitoneal (i.p.) injection in a standard injection volume of 300 µl 30 minutes before the MBT. Mice were divided into the following treatment groups.: 1. *Vehicle (VEH):* 0.9 % saline. 2. *Psilocybin (PSIL)*: 4.4 mg/kg dissolved in vehicle. (This dose was chosen because in a preliminary experiment we found that PSIL 1.5 mg/kg did not significantly affect marble-burying in ICR mice) 3. *Escitalopram (ESC)*: 5 mg/kg dissolved in vehicle. 4. *8-OH-DPAT (DPAT)*: 2 mg/kg dissolved in vehicle containing 5% DMSO. 5. *M100907*: 2 mg/kg dissolved in vehicle containing 5% DMSO. 6. *Buspirone (BUSP)*: 5 mg/kg dissolved in vehicle. 7. *WAY100635 (WAY)*: 2 mg/kg dissolved in vehicle. 8. *M100907 + PSIL* 9. *DPAT + PSIL*. 9. *WAY + PSIL*. 10. *BUSP + PSIL*.

### Marble-burying test (MBT)

MBT (see Supplementary Video) was performed in transparent cages containing ~4.5 cm sawdust, as described by Odland et.al, (2021a). Twenty glass marbles were placed equidistant from each other in a 5 × 4 pattern. The experiment was done under dim light in a quiet room to reduce the influence of anxiety on behavior. The mice were left in the cage with the marbles for a 30-min period after which the test was terminated by removing the mice. Number of buried marbles was counted after 10, 20 and 30 minutes. All mice underwent a pretest without any injection and the number of marbles buried was counted. Only mice that buried at least 15 marbles were selected to perform the test after drug administration. 80% of pretested mice fulfilled this criterion and were used in the definitive experiment which took place at least a week following the pretest.

### Open Field Test (OFT)

The OFT was performed immediately after the MBT to evaluate the effects of the drug treatments on locomotor activity. The apparatus consisted of a square wooden arena (50 × 50 × 40 cm) with white walls and a black floor. Mice were placed individually in the center of the open field and allowed to freely explore the apparatus for 30 min. A camera was used to monitor movement. Total distance travelled (centimeters) was measured by the Ethovision Video Tracking System (Noldus Information Technology BV). After each test, the arena was cleaned with 70 % alcohol solution.

### Head Twitch Response

Head twitch response (HTR) was measured over 20 minutes by means of a magnetometer apparatus as described by Revenga et al (2020) and Shahar et al (Submitted). Briefly, small neodymium magnets (N50, 3mm diameter × 1mm height, 50 mg), were attached to the outer ears of mice. After a 5–7-day recovery period, the ear-tagged animals were placed inside a magnetometer apparatus (supplied by Mario de la Fuente Revenga PhD. of Virginia Commonwealth University) immediately after injection of vehicle or psilocybin 4.4 mg/kg. The output was amplified (Pyle PP444 phono amplifier) and recorded at 1000 Hz using a NI USB-6001 (National Instruments, US) data acquisition system (Revenga et al., 2020). Recordings were performed using a MATLAB driver (MathWorks, US, R2021a version, along with the NI myDAQ support package) with the corresponding National Instruments support package for further processing. A custom MATLAB script was used to record the processed signal which was presented as graphs showing the change in current as peaks (mAh). A custom graphic user interface created in our laboratory was used to further process the recording into an Excel spreadsheet.

### Statistical analysis

The experimental data in all figures are expressed as the mean ± standard error of the mean (SEM). To determine inter group differences, one-, two-, and three-way analysis of variance (ANOVA) were used as indicated. Tukey’s Multiple Comparison Test was used to analyze post-hoc comparisons. p <0.05 (two tailed) was the criterion for significance. Graph Pad Prism, version 9.3.1 software was used for all statistical analyses.

## Results

### Marble-burying Test

Mice administered psilocybin buried 32.84% fewer marbles over 30 minutes than vehicle treated mice (p=0.001, Fig 1a). The effect of psilocybin was not statistically different from the effect exerted by a positive control, the SSRI escitalopram (48.43% reduction in marble-burying relative to vehicle; p<0.0001, Fig 1a). Using a two-way-ANOVA design, we examined the effects of acute treatment with psilocybin and pretreatment with the 5-HT2A antagonist, M100907 (volinanserin) (Fig 1b). A strong main effect of psilocybin was noted (F_1,40_=24.8, p<0.0001). We also observed a main effect of M100907 (F_1,40_=7.7, p=0.008, Fig. 1b). However, there was no interaction between psilocybin and M100907 and the post-hoc comparison of M100907 and vehicle was not significant (p>0.10) while the post-hoc comparison of M100907+psilocybin and vehicle was (p<0.0001). These findings indicate that the mechanism whereby psilocybin reduces marble-burying behavior is likely independent of 5-HT2A receptor signaling.

**Figure 1:**
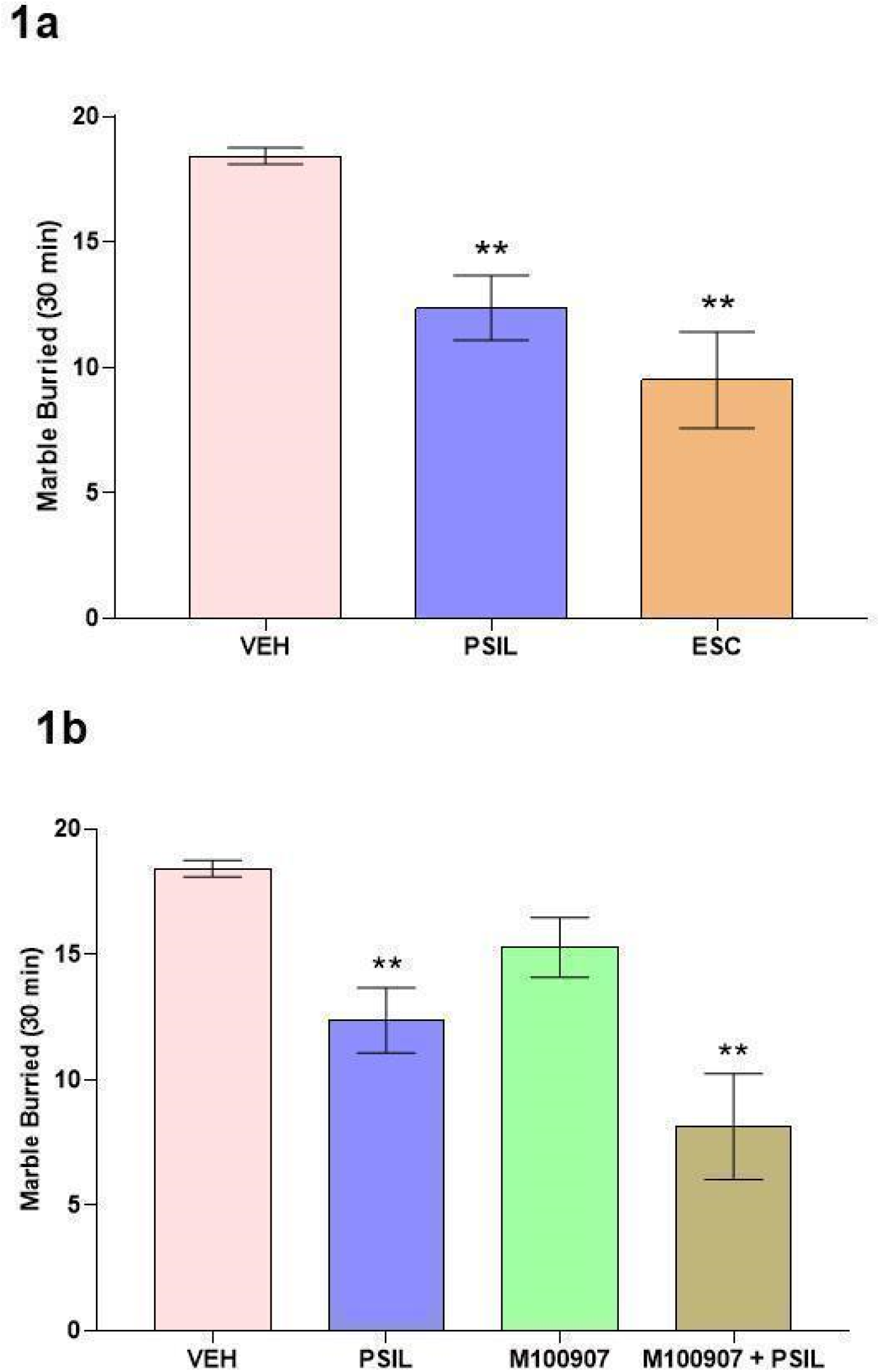
**1a**: Effect of psilocybin 4.4 mg/kg and escitalopram 5 mg/kg on total marbles buried over 30 minutes. One way ANOVA: F_2,35_ = 13.32 p<0.0001. **p<0.01 vs. VEH, n=8-16 (Tukey’s multiple comparisons test). **1b**. Effect of psilocybin 4.4 mg/kg, M100907 2 mg/kg and M100907 2 mg/kg + psilocybin 4.4 mg/kg on total marbles buried over 30 minutes. Two-way ANOVA: M100907 F_1,40_ = 7.74, p=0.008, psilocybin F_1,40_ = 24.80, p<0.0001, Interaction F_1,40_ = 0.169, p=0.68. **p<0.01 vs. vehicle, n=7-16 (Tukey’s multiple comparisons test).

Could stimulation of 5-HT1A receptors underlie psilocybin’s effect on marble-burying? Psilocybin and the 5-HT1A agonist, 8-OH-DPAT, both exerted significant main effects to reduce marble-burying (F_1,37_=10.4, p=0.0026 and F_1,37_=74.3, p<0.0001, respectively, Fig. 2a). However, psilocybin did not interact with 8-OH-DPAT (F_1,37_=0.9, p=0.34 for psilocybin-DPAT interaction term) indicating that 5-HT1A stimulation was unlikely to account for psilocybin’s effect in the marble-burying paradigm. Moreover, the combined effect of psilocybin and 8-OH-DPAT was significantly greater than that of vehicle or psilocybin alone (p<0.0001 and p<0.0001 respectively). To consolidate this observation, we tested the effects of treatment with psilocybin preceded by the 5-HT1A receptor antagonist WAY100635. Using a two-way-ANOVA design, we noted a strong main effect of psilocybin (F_1,61_=42.47, p<0.0001, Fig. 2b), while the main effect of WAY100635 and the psilocybin-WAY100635 interaction were both not significant (F_1,61_=0.4, p=0.521 and F_1,61_=0.0003, p=0.985, respectively, Fig. 2b). Post-hoc testing confirmed that pretreatment with WAY100635 does not block the effect of psilocybin to reduce marble-burying (p>0.10, Fig 2b).

**Figure 2:**
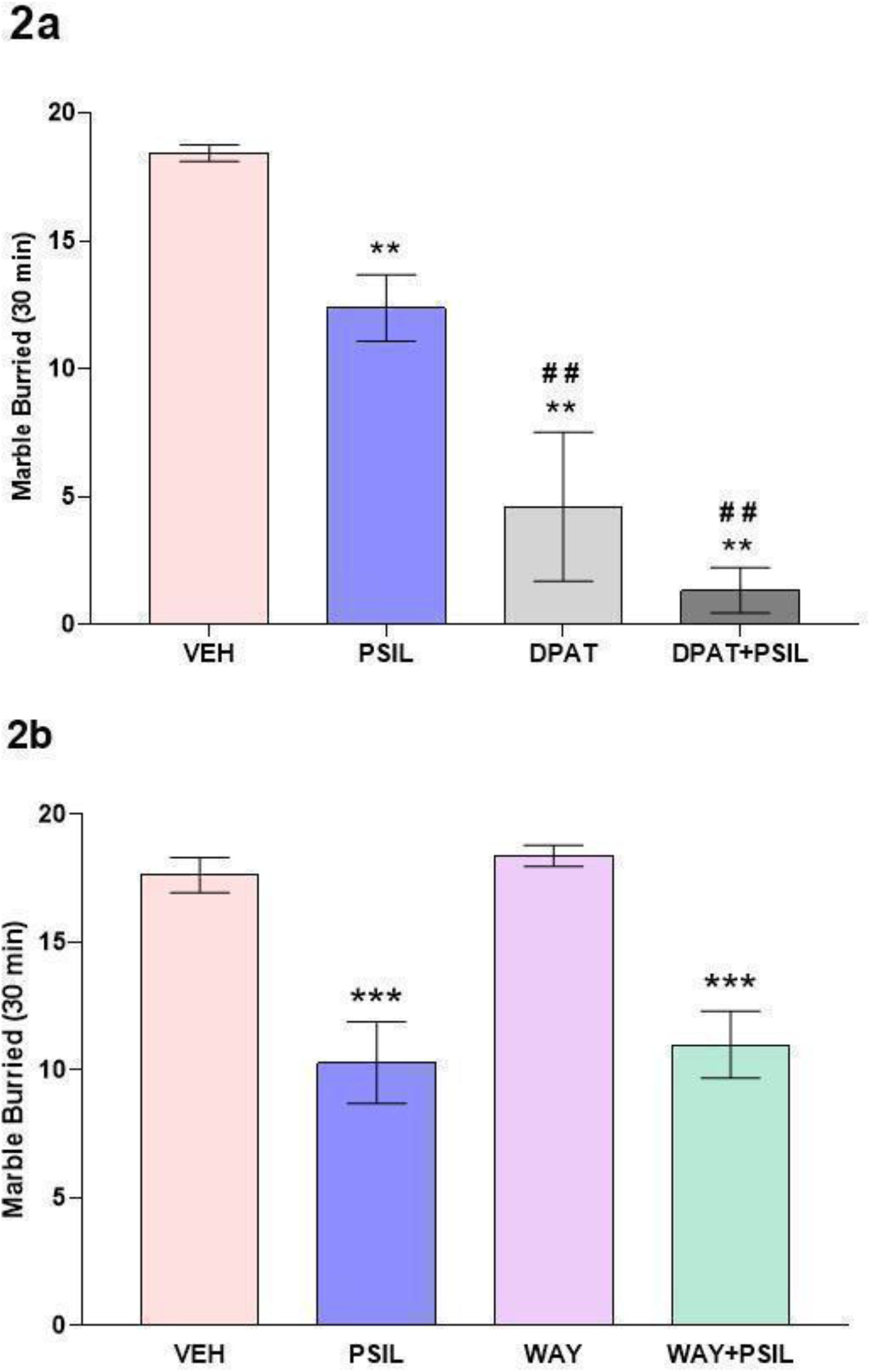
**2a**: Effect of psilocybin 4.4 mg/kg, 8-OH-DPAT 2mg/kg and 8-OH-DPAT 2 mg/kg + psilocybin 4.4 mg/kg on total marbles buried over 30 minutes.Two way ANOVA: psilocybin F_1,37_ = 10.43, p=0.0026, 8-OHDPAT F_1,37_ = 74.25, p<0.0001. Interaction F_1,37_ = 0.9324, p = 0.3405) **p<0.01 vs. VEH. ^##^p<.01 vs. psilocybin, n = 6-16 (Tukey’s multiple comparisons test). **2b**: Effect of psilocybin 4.4 mg/kg, WAY100635 2 mg/kg and WAY100635 2 mg/kg + psilocybin 4.4 mg/kg on total marbles buried over 30 minutes. Two-way ANOVA: WAY100635 F_1,61_ = 0.4162, p=0.5212, psilocybin F_1,61_ = 42.47, p<0.0001, Interaction F_1,61_ = 0.0003, p=0.9845. **p<0.001 vs. vehicle, n=16-17 (Tukey’s multiple comparisons test).

Buspirone is a 5-HT1A receptor partial agonist and a weak dopamine D2 receptor antagonist (Di Ciano et al., 2017; Loane and Politis, 2012). Unlike 8-OH-DPAT, we found an interaction of buspirone with psilocybin (F_1,75_=5.805, p=0.018, Fig. 3a). Thus, although psilocybin and buspirone alone both significantly reduced marble-burying compared to vehicle (F_1,75_=6.53, p=0.015; F_1,75_=68.53, p=0.005; respectively, Fig. 3a), their co-administration did not yield a further reduction in marble-burying. We also tested the effect of pretreatment with WAY100635 on the reduction in marble-burying induced by buspirone (buspirone F_1,59_=19.45, p<0.0001, WAY100635 F_1,59_=3.07, p=0.08, WAY100635 X buspirone F_1,59_=1.2, p=0.274, Fig. 3b). Contrasting with the lack of effect of WAY100635 on the psilocybin-induced reduction in marble-burying (Fig 2b), we found a significant effect of WAY100635 to attenuate the effect of buspirone on marble-burying (buspirone vs. vehicle, p<0.001; WAY100635 +buspirone vs. vehicle, p>0.10).

**Figure 3:**
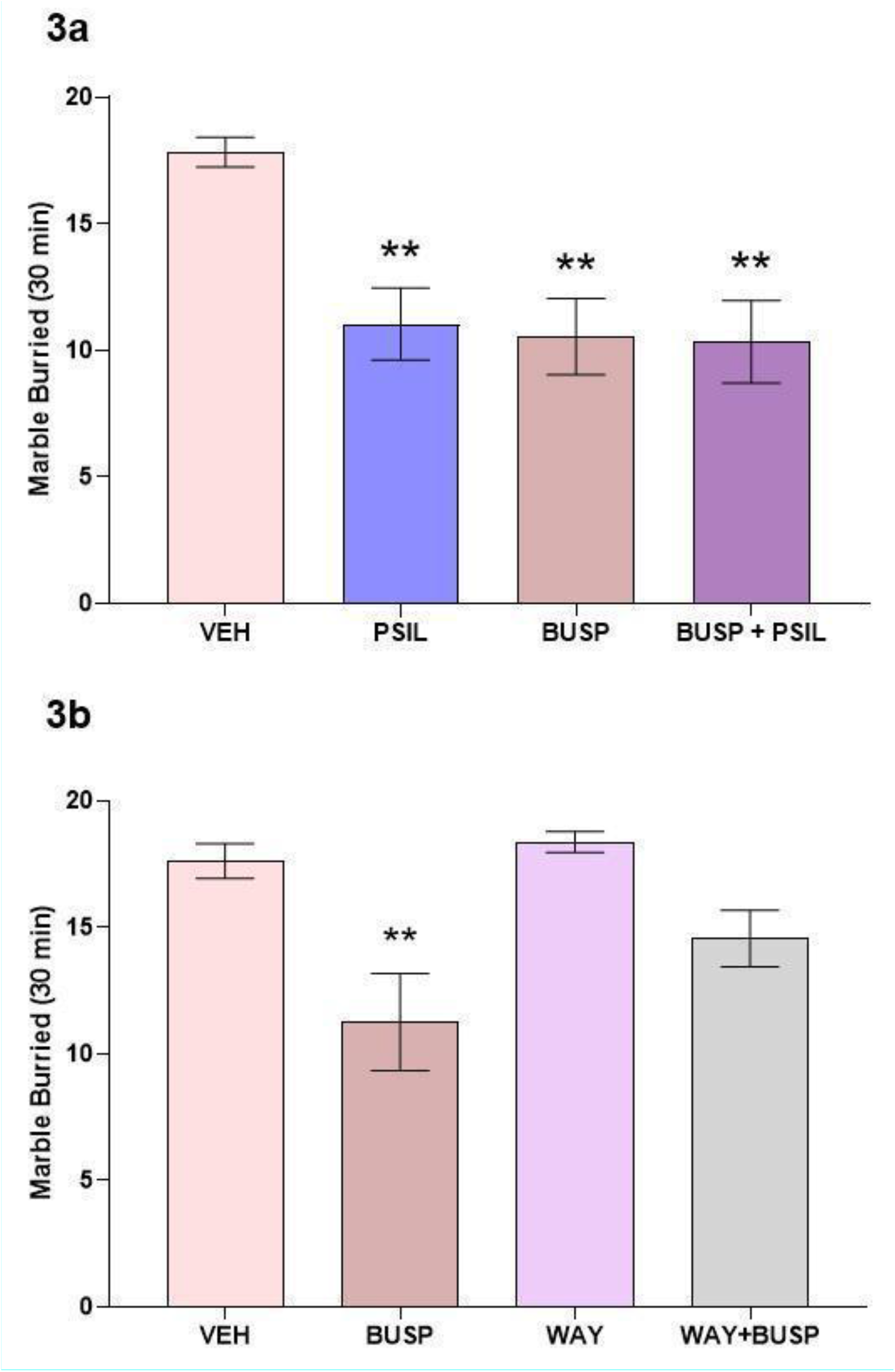
**3a**: Effect of psilocybin 4.4 mg/kg, buspirone 5 mg/kg and buspirone 5 mg/kg+ psilocybin 4.4 mg/kg on total marbles buried over 30 minutes. Two-way ANOVA: buspirone F_1,75_ = 8.532, p=0.0046; psilocybin F_1,75_ = 6.530, p= 0.0126; Interaction F_1,75_ = 5.805, p = 0.0184 **p<0.01 vs. VEH, n=19-20 (Tukey’s multiple comparisons test). 3**b:** Effect of buspirone 5 mg/kg, WAY100635 2 mg/kg and WAY100635 2 mg/kg + buspirone 5 mg/kg on total marbles buried over 30 minutes. Two-way ANOVA: WAY100635 F_1,59_ = 3.078, p=0.084, buspirone F_1,59_ = 19.45, p<0.0001, Interaction F_1,59_ = 1.219, p =0.274. **p<0.001 vs. VEH, n=15-16 (Tukey’s multiple comparisons test).

A key question not addressed in studies thus far is whether the effect of psilocybin and other psychedelic compounds on marble-burying is transient or persistent. We examined marble-burying in a subset of mice treated with vehicle or psilocybin 7 days following the initial MBT. No significant effect of psilocybin was observed (vehicle 18.6 ± 1.6, n=5; psilocybin 17.75± 2.21 n=4; p =0.53). We further examined whether the effect of psilocybin on marble-burying requires a bolus injection of the full dose of the drug (at 4.4 mg/kg) or whether the same effect can be achieved by administering the same quantity of drug in staggered fashion over a period of 3.5 hours i.e. i.p. injections of 1.1 mg.kg every 60 minutes with the MBT performed 30 minutes after the last injection. When administered in this fashion, no significant effect of psilocybin on marble-burying was observed (vehicle 19±0.89 n=6; psilocybin 19±1.32 n=9; p>0.10).

### Open Field Test

Mice were placed in an open field on completion of the MBT and were monitored for 30 minutes using the Ethovision Video Tracking System (Noldus Information Technology BV). As shown in Supplementary Fig. 1a there was no significant difference in distance travelled between vehicle treated mice and those administered psilocybin or buspirone. Similarly, there was no difference in time spent by the mice in the center of the open field (center duration) (Supplementary Fig 1b) or in the periphery of the open field (periphery duration) (Supplementary Fig 1c) under treatment with psilocybin or buspirone compared to treatment with vehicle.

### Evaluation of HTR

To determine whether the dose of psilocybin that inhibited marble-burying in this study would induce a significant increase in the number of head twitches observed in ICR mice as compared to vehicle, HTR was measured in a magnetometer device as described above. We further examined the effect of co-administration of buspirone 5 mg/kg with psilocybin 4.4 mg/kg to determine whether co-administration of buspirone would attenuate the HTR-enhancing effect of psilocybin. Fig 4a shows the time course of the effect of psilocybin and buspirone on HTR. Three way ANOVA with repeated measures showed a significant within-subjects effect of time (F_9,288_ = 5.001, p <0.0032), reflecting the changes in HTR rate during the course of the test; a time by psilocybin interaction (F_9,288_ = 3.224, p = 0.001), reflecting differential, psilocybin-dependent changes in the HTR rate during the course of the test and a triple, time by psilocybin by buspirone interaction (F_9,288_ = 2.687, p=0.0072), reflecting differential psilocybin- and buspirone-dependent, changes in the HTR rate over time. Significant between-subject effects of psilocybin (F_1,32_ = 19.22, p=0.0001) and buspirone were observed (F_1,32_ = 7.483, p=0.0101) and a significant psilocybin by buspirone interaction (F_1,32_ = 5.237, p=0.0289) indicating effects over time. When evaluating the total number of HTRs during the 20-minute measurement period (Fig, 4b) there were significant effects of psilocybin (F_1,32_ = 19.22, p=0.0001) and buspirone (F_1,32_ = 7.483, p=0.0101) and a significant psilocybin x buspirone interaction (F_1,32_ = 5.237, p=0.0289). In post-hoc tests, there was a significant effect of psilocybin to increase HTR compared to vehicle (p<0.001) while buspirone and buspirone + psilocybin effects were significantly lower than the effect of psilocybin (p=0.0002 and p=0.0009, respectively)

**Figure 4:**
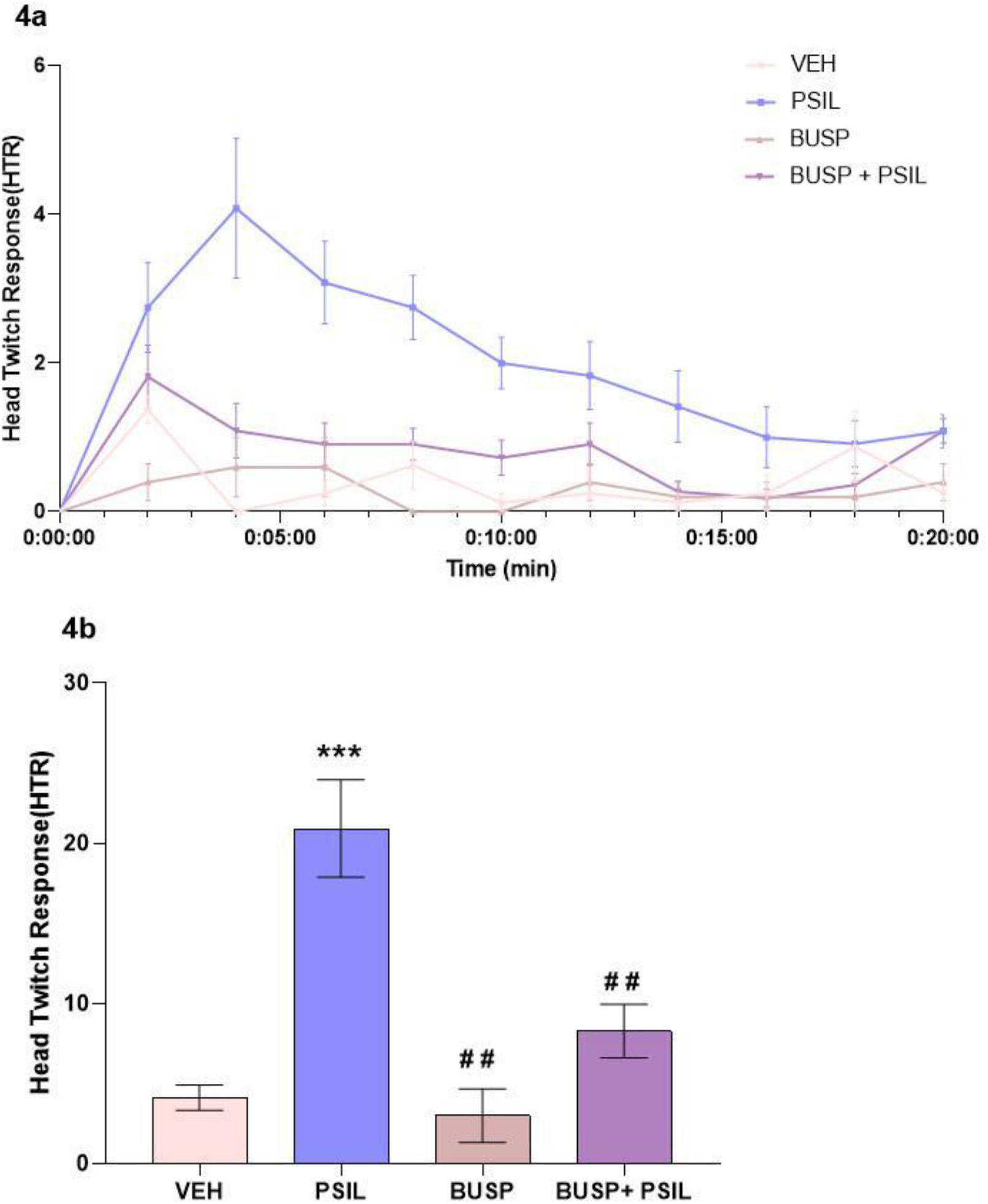
**4a**: Effect of psilocybin 4.4 mg/kg, buspirone 5 mg/kg and psilocybin 4.4 mg/kg + buspirone 5 mg/kg on HTR over a 20-minute measurement period. Three-way ANOVA: Time F _9,288_ = 5.001, p = 0.0032; Time x psilocybin F _9, 288_ = 3.224, p = 0.001; Time x psilocybin x buspirone F _9, 288_ = 2.687, p=0.0072 (within subject effects). psilocybin F _1, 32_ = 19.22, p=0.0001; buspirone F_1, 32_ = 7.483, p=0.0101; psilocybin x buspirone F_1, 32_ = 5.237, p=0.0289 (between subject effects). **4b**: Total HTR over 20 minutes. F_3,32_ = 12.87, p <0.0001; ***p<0.001 vs. vehicle, ^##^p= 0.0002 buspirone vs. psilocybin and p=0.0009 buspirone + psilocybin vs. psilocybin, n=6-12 (Tukey’s multiple comparisons test)

## Discussion

The results of our study support a significant effect of psilocybin to reduce the number of marbles buried in the MBT by male ICR mice when the drug is administered 30 minutes before the test, as previously reported by Matsushima et al (2009) and by Odland et al (2021a). Matsushima et al (2009) studied male ICR mice and observed a significant effect of psilocybin at a dose of 1.5 mg/kg. Odland et al (2021a) studied female NMRI mice and observed a significant effect of psilocybin at 1.0 mg/kg. In our study, 1.5 mg/kg was not sufficient to obtain a significant effect on male ICR mice and we a used dose of 4.4 mg/kg psilocybin for our experiments. The reason for this difference between our study and that of Matsushima et al (2009) is not clear.

As reported by Odland et al (2021a) we found that the 5-HT2A antagonist, M100907 (volinanserin), did not block the effect of psilocybin on marble-burying. In contrast, the anti-marble-burying effect of the 5-HT2A agonist, DOI, could be blocked by M100907 (Odland et al., 2021a). While the reason for this discrepancy is not clear, our data and those of Odland et al. (2021a) are consistent with psilocybin exerting its effects through mechanisms other than 5-HT2A signaling.

Our findings regarding the role of 5-HT1A receptors in the effect of psilocybin on marble-burying are intriguing. As reported by others (Bruins et al., 2008; Egashira et al., 2008), the prototypical 5-HT1A agonist, 8-OH-DPAT, significantly reduced marble-burying in our study. Combined administration of 8-OH-DPAT and psilocybin exerted an additive anti-marble-burying effect. There was no interaction between 8-OH-DPAT and psilocybin, consistent with a differential mode of action exerted by these two compounds. Moreover, pretreatment with the 5-HT1A antagonist, WAY100635, which has been shown to block the effect of 8-OH-DPAT on marble-burying (Egashira et al., 2008) and was shown to have this effect for the 5-HT1A partial agonist, buspirone, in our study, did not attenuate the effect of psilocybin. Taken together, these two findings suggest that the effect of psilocybin to reduce marble-burying is not mediated by the 5-HT1A receptor but by a different, as yet unelucidated mechanism.

While the 5-HT1A partial agonist, buspirone, significantly reduced marble-burying in our study, its effect was not additive to the effect of psilocybin as was the case for 8-OH-DPAT and psilocybin. In this context it is noteworthy that the psychedelic effects induced by psilocybin are attenuated by buspirone as shown by Pokorny et al. (2016) in a study of healthy volunteers. Consistent with this observation we showed that co-administration of buspirone and psilocybin blocked the effect of psilocybin to induce HTR in mice while not affecting its anti-marble-burying action. The clinical implication of this finding is that co-treatment with psilocybin and buspirone could potentially permit the anti-obsessional effects of psilocybin while blocking its psychedelic effects. Further exploration of this interaction is an important topic for further study that has high relevance for the treatment of OCD.

As previously demonstrated by Odland et al (2021a), the effect of psilocybin and buspirone to reduce marble-burying in our study was not a consequence of reduced motor activity. In our measurements of open field activity there was no difference in the effect of psilocybin and buspirone compared to vehicle. Our activity measurements were not performed during the MBT, 30-60 minutes after the psilocybin injection but after the MBT, 60-90 minutes after psilocybin administration. Nevertheless, it is noteworthy that the activity measurements performed by Odland et al (2021a) were during the MBT and no difference in the effect of psilocybin compared to vehicle was observed.

We did not find an effect of psilocybin on marble-burying that extended beyond that observed after 30 minutes. No effect was observed when a subset of mice was retested after 7 days. Also, we found that a bolus injection of psilocybin was needed to achieve an effect on marble-burying and there was no effect of psilocybin on the MBT when the same dose was spaced over 4 hours rather than being administered at once. Implications for the anti-obsessional effect of psilocybin in humans await studies in which the longer-term effects of psilocybin administration on OCD are examined and the dose required is clarified. In the study by Moreno et al (2006) clinical observations were not performed beyond 24 hours following oral administration of psilocybin. However, doses of psilocybin that did not induce prominent psychedelic effects were found to have anti-obsessional effects while in our study a bolus of drug sufficient to induce prominent HTR in mice was required, as demonstrated by our HTR data.

In conclusion, the results of our study confirm the previously reported effect of psilocybin to reduce marble-burying in mice, confirm that this effect is not blocked by a 5-HT2A receptor antagonist and suggest that the effect is not mediated by 5-HT1A receptors since the effects of psilocybin and 8-OH-DPAT are additive and the effect of psilocybin is not blocked by the 5-HT1A antagonist, WAY100635. The results further show that co-treatment with the 5-HT1A partial agonist, buspirone, blocks effects of psilocybin on HTR, a rodent correlate of psychedelic effects while not impeding its effect on marble-burying. Further studies are indicated to identify the receptor mechanisms of psilocybin in reducing marble-burying in mice with implications for the mechanism of the putative anti-obsessional effect of psilocybin in humans.

## Acknowledgments

We thank Mario de la Fuente Revenga PhD, Virginia Commonwealth University, for providing the magnetometer device and for valuable advice.

## Author contributions

TL and BL conceived and designed the study. TL, BL, SS, AB and OS planned the experiments. SS, AB, OS CY, MS and AS performed the experiments, SS, BL, GW and AL performed the statistical analyses. SS, AL, GW, BL and TL wrote the paper. All the authors read and approved the manuscript.

## Supplemental materials

Supplemental Fig. 1

Supplemental Video

