## Supplemental Material for "Effect of psilocybin on marble-burying in ICR mice: Role of 5-HT1A receptors and implications for the treatment of obsessive-compulsive disorder"

**Singh et al.**

**SUPPLEMENTAL MATERIAL**

Supplemental Figure 1

Supplemental Video

**Supplemental Fig. 1**:

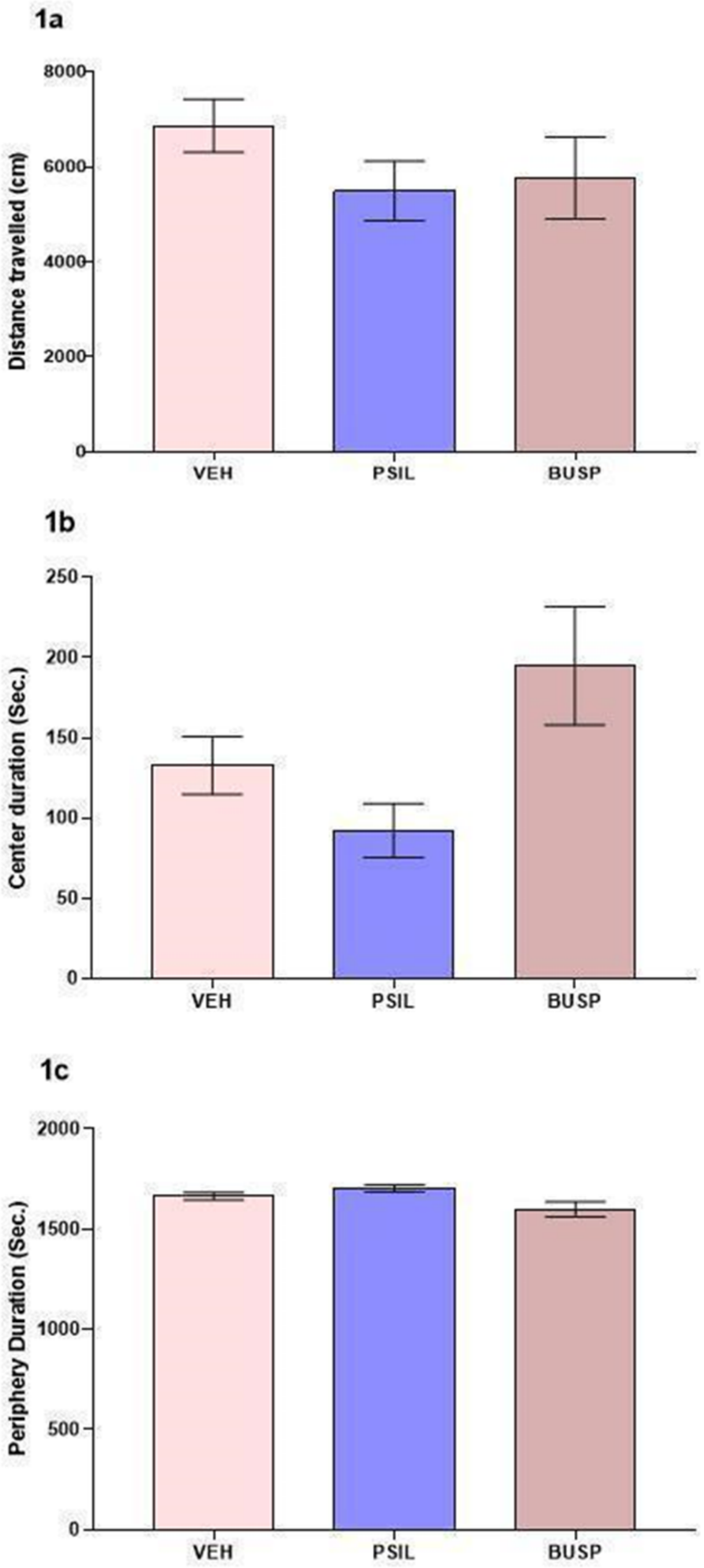

**1a**: Effect of psilocybin 4.4 mg/kg and buspirone 5 mg/kg on distance travelled in the open field over 30 minutes. One way ANOVA: $F_{2,30}$ = 1.044; p=0.3645 . p N.S. vs. VEH

**1b**: Effect of psilocybin 4.4 mg/kg and buspirone 5 mg/kg on time spent in the center of the open field over 30 minutes. One way ANOVA: F 2, 30 = 4.934; p=0.0140. p N.S. vs. VEH.

**1c**: Effect of psilocybin 4.4 mg/kg and buspirone 5 mg/kg on time spent in the periphery of the open field over 10 minutes. One way ANOVA: F 2, 30 = 5.003; p = 0.0133. p N.S vs VEH, n= 9-15 (Tukey's multiple comparisons test).

### SUPPLEMENTAL VIDEO

This video shows ICR mice engaged in marble-burying. This behavior serves as the basis for the marble-burying test. Mice did not receive any pretreatment before being placed in the test cage which contained twenty marbles equidistant from each other in a 5 × 4 pattern. The experiment was done under dim light in a quiet room to reduce the influence of anxiety on behavior. The mice were left in the cage with the marbles for a 30-min period after which the test was terminated by removing the mice. Number of buried marbles was counted after 10, 20 and 30 minutes.

https://drive.google.com/file/d/1n5oKtI4ZyewVwn1suDFXxifdbwjO4Wtt/view?usp=sharing

(Filmed by Dr Alexander Botvinnik, Biological Psychiatry Laboratory and Hadassah BrainLabs, Hadassah Medical Center, Hebrew University, Jerusalem, Israel.)
